# Cofilin Regulates Filopodial Structure and Flexibility in Neuronal Growth Cones

**DOI:** 10.1101/2021.09.17.460569

**Authors:** Ryan Hylton, Jessica Heebner, Michael Grillo, Matthew Swulius

**Affiliations:** Department of Biochemistry and Molecular Biology, Penn State College of Medicine; Hershey, PA, USA

## Abstract

Cofilin is best known for its ability to sever actin filaments, and facilitate cytoskeletal recycling inside of cells. At higher concentrations, *in vitro*, cofilin stabilizes a more flexible, hyper-twisted state of actin known as “cofilactin”, but a structural role for cofilactin, *in situ*, has not been observed. Combining cryo-electron tomography and live-cell imaging in neuronal growth cones, we show that filopodial actin bundles can switch between a fascin-linked and a cofilin-decorated state, composed of hyper-twisted cofilactin filaments. These cofilactin bundles contribute to the flexibility of filopodial actin networks, thus regulating growth cone searching dynamics. Our results provide mechanistic insight into the processes underlying proper brain development, as well as fundamentals of cytoskeletal mechanics inside confined cellular spaces.

## Introduction

In the developing nervous system, functional neural circuits are constructed via a process of chemically and mechanically induced neurite guidance (*1*–*4*). Here, neurites are directed toward their eventual synaptic partners by the coordination of actin polymerization and depolymerization within the “growth cone” found at their distal tips (*1*–*3*, *5*–*9*). When grown in culture, the growth cone is typically fan-shaped with filopodial protrusions at the leading edge, which are connected laterally by a lamellipodial veil made of shorter, branched actin networks (*2*). Both of these structures constitute the growth cone’s peripheral domain, and are highly enriched in filamentous actin (F-actin) (**Fig. 1A**) (*2*, *3*, *5*–*8*). The growth cone advances and turns by integrating both attractive and repulsive cues from the environment and converting them into signaling cascades that drive remodeling of the actin cytoskeleton (*2*, *3*, *10*–*15*). Here, filopodia act as antennae, detecting and responding to extracellular cues, while actin network modification in the lamellipodia moves the membrane and drives the growth cone towards its destination (*2*, *5*, *16*–*18*).

**Fig. 1.**
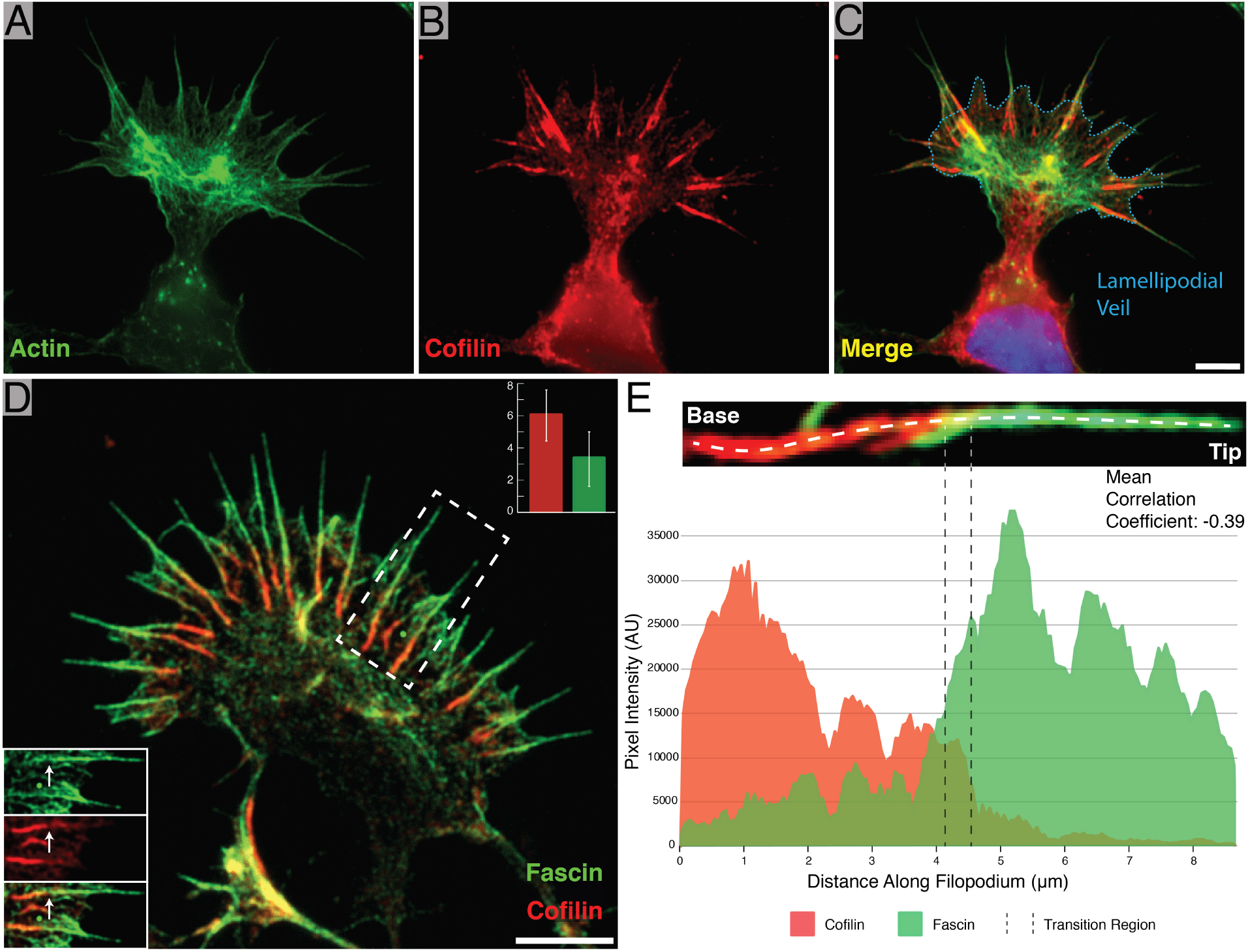
Identification of cofilin-rich regions at the base of neuronal growth cone filopodia. **(A-C)** Immunofluorescence images of phalloidin (Alexa Fluor 488) alone (A), cofilin (Alexa Fluor 594) alone (B), and a merge of the two (C). Most of the cofilin signal emanates from linear aggregates near the base of filopodia. The blue dashed line in (C) marks the position of the lamellipodial veil. **(D)** Merged immunofluorescence image of a growth cone with fascin (Alexa Fluor 488) and cofilin (Alexa Fluor 594) labeled, showing cofilin-rich regions at the base of filopodia as in (C). Bottom-left inset: Split view of the boxed-out region. The white arrow points to the same location in each image and shows the point at which the fascin signal drops off and the cofilin signal intensifies. Top-right inset: Lengths of fascin-rich filopodial region in filopodia with (red) and without (green) a cofilin-rich base (error bars represent S.D.). **(E)** Representative line scan intensity profile of a single filopodium showing the distribution of fascin and cofilin. The image above the graph shows a close-up view of the measured filopodium and the location where the line profile was drawn. The transition region is marked by two dashed lines. Scale bars: (C) = 5 μm (this also corresponds to A, and B, (D) = 5 μm.

This motion is largely driven by polymerization of actin at the cell’s leading edge where the barbed ends of actin are concentrated (*5*), and it manifests in live-cell movies as a retrograde flow of actin from the leading edge toward the back of the lamellipodium (*5*). Retrograde flow is seen both in filopodial and lamellipodial networks, and is regulated through interactions with the myosin family of motor proteins, which increase flow rates (*19*, *20*). It was also shown that filopodial extension is inversely correlated with retrograde flow (*19*).

Remodeling of filopodial actin networks is facilitated by an array of actin binding proteins (*6*, *21*). For example, the actin cross-linking protein fascin connects actin into filopodia made of hexagonally-packed, linear F-actin with regular lateral spacing (∼12.3 nm) (*22*–*27*). The most widely described function for cofilin, on the other hand, is to sever F-actin (*28*–*31*) and regulate its polymerization/depolymerization kinetics (*28*, *31*, *32*). Previous work has shown that cofilin is necessary for normal neurite outgrowth (*6*, *33*, *34*) and that outgrowth can be increased by its overexpression (*35*).

Multiple groups have demonstrated that purified cofilin can bind to F-actin at a 1:1 molar ratio (*36*, *37*), and induce a shortened helical pitch in the filament (∼27 nm crossover length compared to ∼37 nm for normal F-actin) (*37*–*39*). These cofilin-saturated actin filaments (known as cofilactin) can be stabilized at higher concentrations of cofilin and studied *in vitro* (*37*, *38*, *40*), but it is still not known whether stable cofilactin filaments play a functional role *in situ*. Light microscopy and fluorescence anisotropy of *in vitro* filaments has shown that cofilactin is more compliant in both bending (*41*) and twisting (*42*) compared to bare actin, suggesting it could alter the mechanical properties of actin networks.

It is not clear how this filament hyper-twisting by cofilin affects fascin-linked bundles of actin, or how those interactions influence growth cone dynamics. Previous research in non-neuronal cells suggests that cofilin works synergistically with fascin at the tips of filopodia to sever actin filaments during filopodial retraction (*43*). In this model, the twisting cofilactin filaments in the tips of filopodia generate tension against the fascin cross-links, and these localized stress-points increase severing events. Conclusions are mixed *in vitro*, where some studies show increased filament severing by cofilin in the presence of cross-linkers (*44*), while others show different effects (*45*, *46*).

Here we reveal, in neuronal growth cones, that filopodial actin bundles switch between a fascin-linked and a cofilin-decorated state, and that this switch alters the structure of filopodial actin networks to exclude fascin cross-linkers. Additionally, filopodial bending occurs primarily at the transition region between the cofilin-rich base and the fascin-rich tip. Our data, combined with findings from the literature, suggest that this structural transition regulates filopodial mobility and, ultimately, neurite outgrowth, by tuning the flexibility of actively searching filopodia as well as regulating interactions with motor proteins, such as myosin II (*20*, *47*, *48*).

## Results

### Cofilin decorates the base of growth cone filopodia and forms an inverse gradient with fascin

We sought to examine the impact of cofilin on the structure and flexibility of actin filament networks in growth cone filopodia, and we started by observing the distribution of cofilin within rat hippocampal growth cones using phalloidin and immunofluorescence (IF) labeling. We observed that in filopodia, F-actin runs from the tip of the protrusion to within the growth cone body, internal to the lamellipodial veil. On approximately half of these filopodial bundles (46%, n = 1,049 filopodia) cofilin labeling was brightly distributed along the last third of the filopodial bundle length (35.3% +/− 1.5 S.E.M., n = 69 filopodia), nearest to the base (**Fig. 1A-C**). We did not observe significant cofilin staining at the tips of growth cone filopodia, such as that seen during filopodial retraction in non-neuronal cells (*43*), but the cofilin-rich region is sometimes seen extending out into the protrusions. One possibility is that the immunostaining protocol is to blame for this discrepancy, given that cofilin-rich bundles are easily disrupted by certain permeabilization protocols, as we determined for this study (**Fig. S1**). It’s also possible that cofilin has cell type-specific functions, which seems likely given the nature of their evidence.

Growth cones double-labeled for fascin and cofilin revealed an inverse gradient of these two proteins running the length of the filopodia, with fascin enriched near the tip, cofilin enriched near the base, and a transition region between them (further defined in the Supplementary Text), where colocalization occurs (Pearson’s Correlation Coefficient = 0.39 +/− 0.06 S.E.M., n = 26 filopodia, **Fig. 1D & E**). Figure S2 shows representative scattergrams of a whole filopodium and of a transition region. These data together suggest that fascin is largely excluded from the cofilin-rich base of growth cone filopodia, except for in the region where the two portions of the bundle intersect. Finally, measuring the length of filopodia both with (n = 97) and without (n = 72) cofilin at their base revealed that, on average, the fascin-rich regions of filopodia were twice as long when a cofilin-rich base was present (6.1 μm +/− 1.6 S.D. vs. 3.3 μm +/− 1.6, respectively, **Fig. 1D**), suggesting a role in the regulation of filopodial length.

### Cofilin hyper-twists actin within filopodial bundles and rearranges filament packing

Cryo-ET was used to examine the structure of actin bundles along filopodia, from their tip to their base within the body of the growth cone (**Fig. 2A**). We found that their distal tips typically contained hexagonally-bundled, cross-linked, linear actin filaments, as previously described in other filopodia (*22*, *25*) (**Fig. 2B, Movie S1**). Closer to the base of the filopodia we often found hexagonally-packed bundles of what appeared to be cofilactin filaments, each with a reduced helical pitch (27.95 nm +/− 0.21 S.E.M, n = 56 twists, **Fig. 2C, Movie S2**). The shortened twist of these filaments matched previous reports of cofilactin filament structure determined *in vitro* (*36*, *37*).

**Fig. 2.**
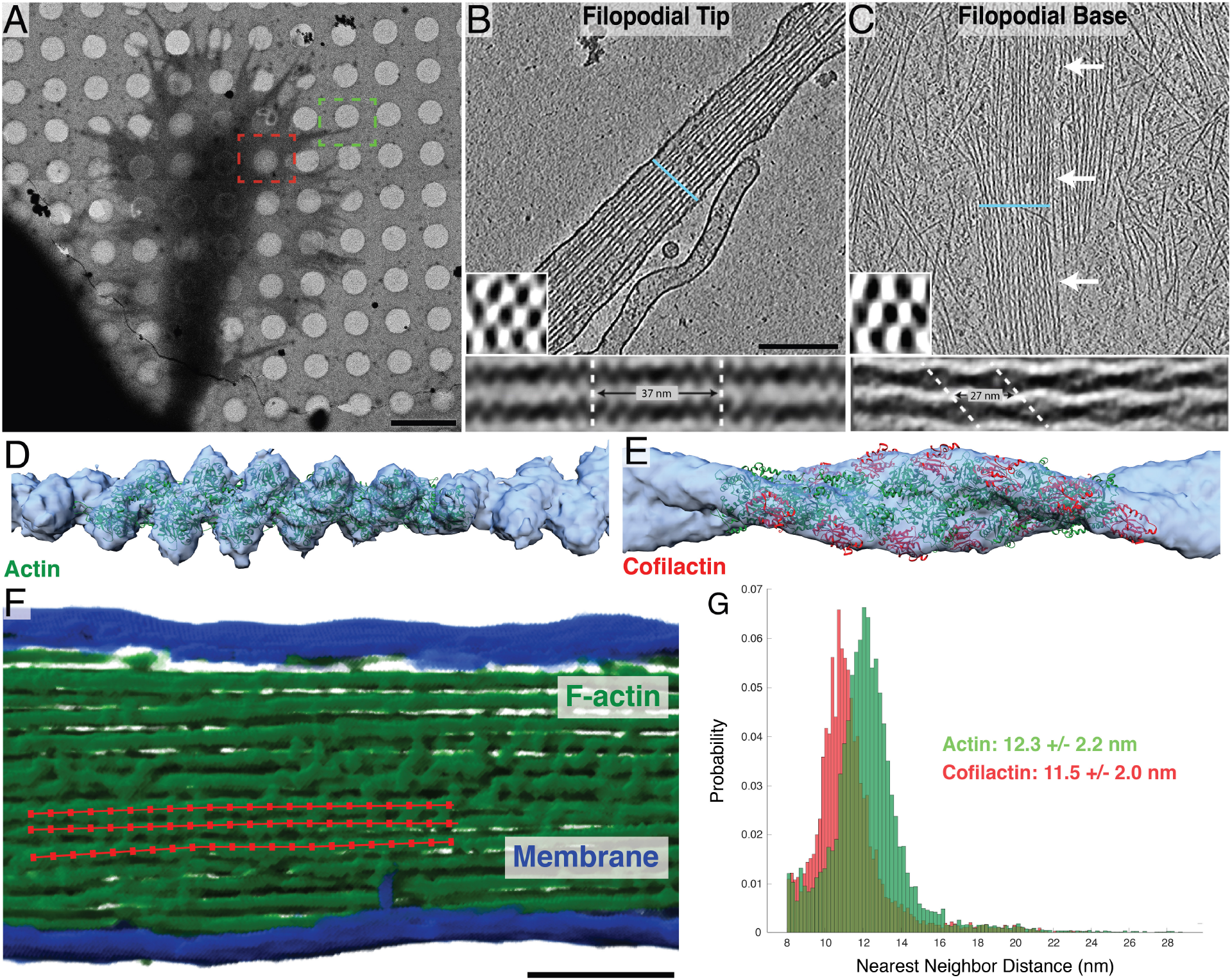
Structural features of neuronal growth cone filopodia and their associated cofilactin bundles. **(A)** Overview image of a cryo-preserved growth cone on a Quantifoil EM grid. The green and red boxes represent growth cone regions similar to where the tomograms in (B) and (C) were imaged, respectively. **(B & C)** 5 nm-thick slices of a tomogram from the tip (B) and base (C) of growth cone filopodia. In (B), a bundle of actin filaments fills the entire cytoplasm. In (C), branched networks of individual actin filaments can be seen surrounding a central bundle of hyper-twisted cofilactin filaments. White arrows point to the bundle. Lower-left insets: 68 nm-thick transverse cross-sections through each bundle, illustrating the hexagonal packing of filaments. The blue line in the main images show the plane from which the insets were taken. Bottom insets: Subtomogram averages of filament pairs in filopodial tips (below (B)) or in cofilactin bundles at the filopodial base (below (C)). Cofilactin filaments have a shorter helical twist than F-actin and are out of phase with adjacent filaments. **(D)** EM map (blue) resulting from the subtomogram averaging of actin filaments in filopodial tips, and rigid body fitting of a previously reported atomic structure for F-actin (PDB ID: 3G37; green). **(E)** EM map (blue) resulting from the subtomogram averaging of cofilactin filaments near the base of filopodia, and rigid body fitting of a previously reported atomic structure of cofilactin (PDB ID: 3J0S; actin is green and cofilin is red). **(F)** Segmented filopodial protrusion with a schematic of filament centerlines overlaid (red). These lines are comprised of a series of points that were used for nearest neighbor analysis. **(G)** Nearest neighbor histograms showing the cumulative total of three normal actin filopodial bundles (green) and three cofilactin bundles (red). Scale bars: (A) 5 μm, (B) 200 nm, this also corresponds to the image in (C), (F) 100 nm.

Averaging of filament pairs within each region showed that neighboring filaments in the tip run parallel to one another, and that their helical pitches are in phase (**Fig. 2B, bottom inset**). Nearer to the base, however, filaments within cofilactin bundles are 90° out of phase with one another, such that the thick portion of one filament is adjacent to the thin portion of its neighbor (**Fig. 2C, bottom inset**). To further confirm that these hyper-twisted filaments were indeed cofilactin, subtomogram averages of fascin-linked actin and cofilactin within different bundles were generated, and fit with their corresponding atomic models (F-actin, PDB ID: 3G37 and cofilactin, PDB ID: 3J0S) (**Fig. 2D & E**)

Next, we measured the interfilament distance within fascin-linked (12.3 nm, n = 9,397) and cofilactin bundles (11.5 nm, n = 7,117), and found that cofilactin filaments are packed 0.8 nm closer to each other (**Fig. 2F & G, Fig. S3).** This reduced interfilament distance further supports that fascin is not the primary cross-linking molecule in cofilactin bundles. Taken together with our IF results (**Fig. 1D & E**), it appears that the structural transition to cofilactin bundles excludes fascin from filopodial bundles.

### Cofilin distribution regulates the flexibility of whole filopodia

To characterize the function of cofilin-rich bundles in the growth cone, we collected super-resolution movies (Zeiss Airyscan 2) of rat hippocampal neurons co-expressing fluorescently labeled cofilin and LifeAct (a small peptide that binds F-actin; (*49*)) (**Fig. 3A, Movie S3**). We observed two basic phenotypes among filopodia in our movies, which we defined as “resting” and “searching” (see Methods for detailed description and **Fig. S4**). In resting filopodia, the cofilin-rich base remained behind the lamellipodial veil, and the bulk of filopodial movement is seen as lateral translations near the base, which creates a smear in the maximum intensity projection (MIP) of our movie frames (**Fig. 3B**). In the searching phenotype, however, lateral movement of the tip, due to filopodial bending, was primarily seen (**Fig. 3C**).

**Fig. 3.**
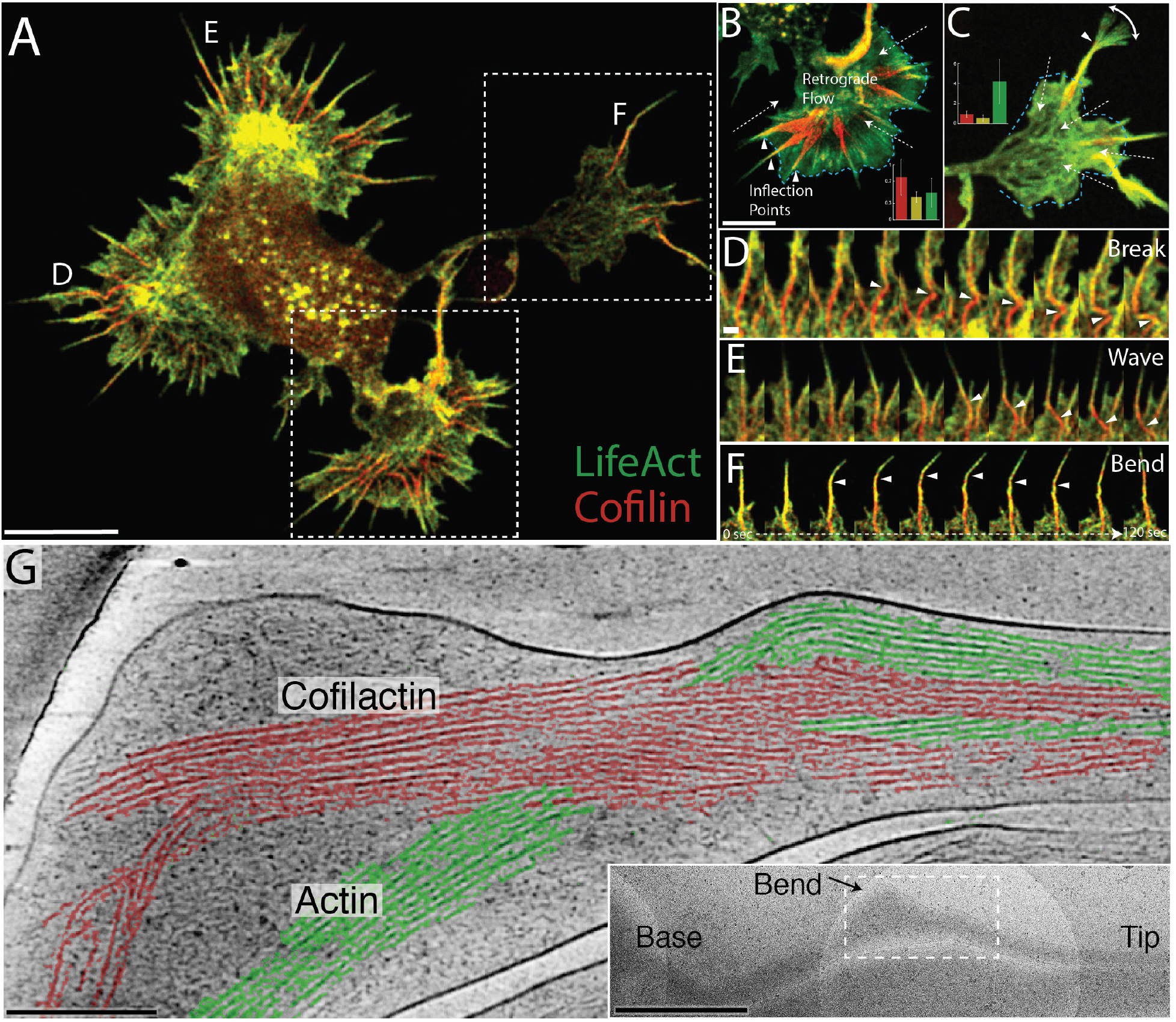
Cofilactin bundles facilitate bending and breaking of filopodial protrusions. **(A)** Single image of a whole cell expressing tdTomato-LifeAct (pseudocolored green) and EGFP-cofilin (pseudocolored red). **(B and C)** Maximum intensity projections (MIP) of lower and upper boxed regions in (A), respectively. MIPs includes 40 images at three-second intervals for 2 minutes total. Filopodia in (B) are examples of “resting” filopodia and (C) shows a “searching” filopodium. Dashed arrows indicate the direction of actin retrograde flow., and arrow heads indicate inflection points seen along the flexing filopodial bundle. The blue dashed line marks the position of the lamellipodial veil. Graphs in (A) and (B) show the average width of cofilin (red), actin (green), and the inflection points (yellow) within MIPS (n = 64 resting and 15 searching filopodia. The y-axis values are in μm. Error bars represent S.D. **(D-F)** 2-minute movie montages showing behaviors exhibited by cofilin-rich filopodia in those designated with the corresponding label in (A). **(G)** 8 nm-thick tomographic-slice through a prospective transition region where actin is segmented in green and cofilactin in red. In this case, both bundle types coexist and interact at a bend in the filopodium. Inset: TEM overview image showing the location of the tomogram in the main panel. Scale bars: (A & B) 5 μm (scale bar in (B) also corresponds to image in (C), (D) 500 nm (also corresponds to (E and F), (G) 200 nm, inset in (G) 1 μm.

To classify filopodia into these two groups we measured the width of the cofilin-rich base, the inflection point, and the LifeAct-rich tip within MIPs made from 2-minute intervals of our movies (**Fig. S4A)**. As expected, measurements from resting and searching filopodia revealed an inverse relationship between the maximum widths of cofilin- and LifeAct-rich regions (**Fig. 3B & C, inset graphs**), verifying that the base is more mobile in resting filopodia and the tip is more mobile in searching filopodia. In both cases, however, the inflection points along filopodia were precisely at the transition region, where colocalization of F-actin and cofilin is highest (**Fig. 3B**), or where the gradient across them was most abrupt (**Fig. 3C, Fig S4**). Frame-by-frame inspection of our movies showed that movement in resting filopodia was largely produced by lateral shifts and breaking within cofilin-rich filopodial bundles (**Fig. 3D, Movie S4**), whereas the movement of searching filopodia produced phenomena such as waves in the base and bending toward the tip (**Fig. 3E & F, Movie S5 & 6**). Of 79 filopodia examined, 64 (81%) were resting and 15 (19%) were searching.

The cofilactin bundles in our tomograms of wildtype growth cones were distributed from ∼8 microns behind the lamellipodial veil to ∼7 microns beyond the veil, within the membrane-bound protrusion (n = 14). If any of our tomograms captured transition regions, we expected to see a mixture of fascin-linked filaments and bundles of cofilactin. We found multiple candidates, especially where we saw curvature/bending in filopodia (**Fig. 3G & S5, Movie S7**).

The mechanism of bending is not entirely clear. However, based on our current tomograms, our immunofluorescence colocalization data, and live-cell movies of LifeAct-to-cofilin transition regions, we favor a model where an inflection point is created between the stiffer, fascin-linked tip of the filopodium and its more flexible cofilactin base. We hypothesize that the location of the inflection point along a filopodium is determined by the balance between actin polymerization and myosin-driven retrograde flow, and that it represents the region of the bundle with the greatest proportion of actin-to-cofilactin boundaries. This model also fits with data showing that individual filaments tend to bend and break at cofilactin-to-actin boundary regions (*50*), as well as cryo-EM data showing cofilin binding disrupts longitudinal interactions between neighboring actin monomers (*51*). Depending on the filopodial state (resting vs. searching) the inflection can be broad, or very abrupt, causing filopodia to bend sharply, suggesting that the length of the hinge point helps to determine the angular range of filopodial tip movement (**Fig. 3B & C**). We were not able to resolve the details of these transition regions within the bundles in our tomograms, but in one of our examples (**Fig. 3G**) a bundle is split into two halves, one made of actin and one made of cofilactin. In another tomogram, however, a more abrupt transition region is present at a bend in the filopodium, but the details of the transition region are obscured by its bending around the edge of the carbon support (**Fig. S5**).

By examining our tomograms closely, we were able to develop a model for how the structure of filopodial bundles is regulated by cofilin binding (**Fig. 4A)**. It appears that upon cofilin decoration of filopodial actin, the filament pitch is shortened and every other column of filaments rotates around its long axis by 90° with respect to the neighboring column. This produces a hexagonally-packed bundle of alternating filament orientations, with filaments on the same layer running 90° out of phase with one another, as can be seen directly in the raw data (**Fig. 4B**). We hypothesize that the transition to cofilactin bundles causes a reconfiguration/tightening of the bundle that ejects fascin cross-linkers from the filopodia (**Fig. 4C; Fig. S6A & B**), due to steric hinderance, generating functionally distinct regions along the filopodial length.

**Fig. 4.**
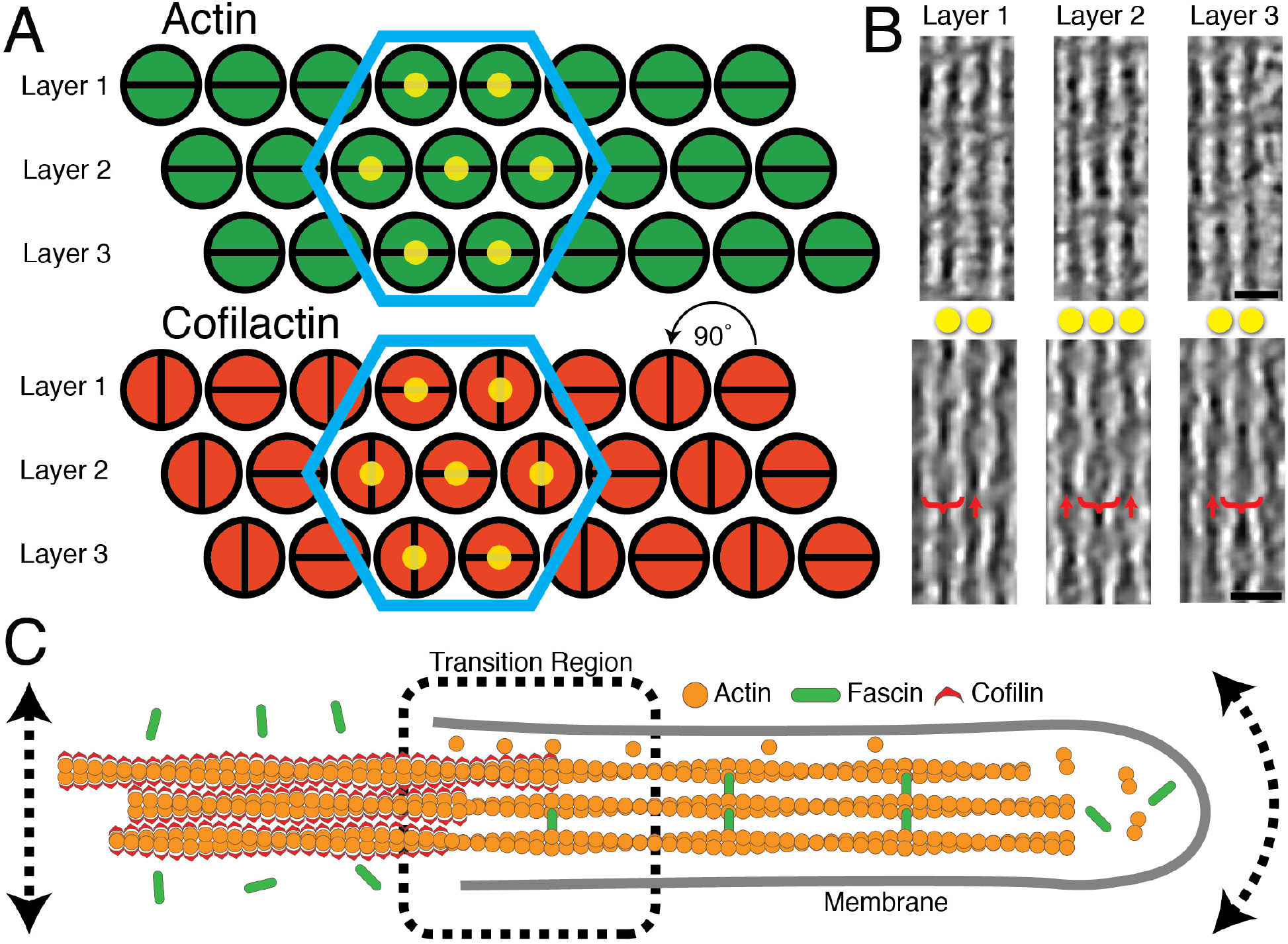
Current Model. **(A)** Schematic model of the higher-order structure of actin (fascin-linked) and cofilactin bundles in growth cone filopodia (as they appear looking down the long axis of the bundle). Filaments in both bundle types are organized in layers and hexagonally packed (blue hexagons). Filaments in the actin bundle are all twisting in phase with one another, but in the cofilactin bundle filaments are rotated 90° with respect to their neighbors in the same layer. This creates columns of filaments that are oriented similarly to one another. The yellow dots on filaments in the hexagon correspond to filaments in (B). **(B)** 17 nm-thick slices through tomograms of a filopodial tip (top) and a cofilactin bundle (bottom). In the cofilactin bundle, brackets show the wide portion of the helical twist while red arrows show the thin portion. The yellow dots correspond to the dots in (A). For instance, the bracketed filament in layer two is directly between the two filaments in layers 1 and 3, only on a different Z-plane. **(C)** Summary of findings presented throughout the paper. In our current model, fascin is dislodged from actin filaments in the cofilin-rich region of the filopodia, and these filaments are instead cross-linked through a different mechanism. The location where actin and cofilactin regions meet creates a transition region that acts as a hinge, enhancing lateral and angular movement of the filopodia at the base and tip, respectively. Scale bars in (B) are 20 nm.

We believe that the functional consequence of the switch between these two bundle types is to generate a hinge-point (or perhaps a universal-type joint) between the stiff filopodial protrusion and its more flexible intracellular base (**Fig. 4C**), and that this flexibility tuning is important for facilitating the bending behavior seen in our searching filopodia. This is not only supported by our live-cell and tomography data, but it is also well established in the literature that cofilactin twists and bends more than F-actin (*41*, *42*). Whether these properties scale up to the level of a whole filopodium is not known, but it is rational to think they would. For instance, increased torsional flexibility could account for the alternating filament rotations seen between filaments within cofilactin bundles (**Fig. 4B**, (*42*)), and increased flexibility (*41*) could explain the motion of cofilin-rich bases in resting filopodia (**Fig. 3B**).

## Discussion

In this study, we present the first evidence for the structural conversion of whole fascin-linked filopodial actin bundles into bundles of hyper-twisted cofilactin filaments. This conversion takes place within discrete transition regions along the length of filopodia that can be observed in our movies of whole growth cones undergoing retrograde flow. The variable location of this transition region (from 2.6 microns behind the lamellipodial veil to 2 microns beyond, as measured in our MIPs, n = 79) suggests it functions as a tunable hinge-point, allowing it to move from behind the lamellipodial veil and into the filopodial protrusion, where it facilitates the searching behavior seen in the tips of a subset of filopodia (**Fig. 3C**).

There are still more open questions, however. For instance, what cross-links cofilactin bundles? Presumably something shorter than fascin, since cofilactin bundles are more tightly packed (**Fig. 2G**), but immunolabeling for other actin cross-linkers such as fimbrin, alpha-actinin and palladin did not show strong staining in our filopodia (data not shown). It’s possible that cofilin itself cross-links bundles through self-associative properties (*52*, *53*). When cofilactin filaments are arranged according to our model, there is potential for interaction between cofilins on neighboring filaments at the interfaces where their helical twists are in phase with one another (**Fig. S6C-E**). In our model, this in-phase boundary only occurs between filaments on different layers of the bundle, and, at the closest points, neighboring cofilin atoms are ∼2 nm apart. This is too far apart for bonds to form regularly, but based on the range of distances measured by our nearest neighbor analysis (**Fig. 2G**), it is possible that a subset of cofilins are close enough to touch at random locations within the bundle. Another possibility is that cofilin could oligomerize to span this distance. Indeed, cofilin oligomers have been shown to bundle actin *in vitro* while monomers do not (*53*).

Little is known about the structure of cofilin oligomers, but cysteines 39 and 147 have been implicated in the formation of inter-cofilin disulfide bonds that drive the formation of “cofilin/actin rods”, which are rod-like bundles of actin and cofilin that form in neurons under oxidative stress (*54*–*56*). While a subset of these rods persist after the removal of stress, most of them disappear in a reversible process (*57*). Based on this, we thought it possible that cofilactin bundles in growth cone filopodia are cross-linked by the same mechanism. If so, it is unlikely a direct cofilin-to-cofilin bond since the closest pair of cofilins on neighboring filaments puts these residues ∼5 nm from each other. Interestingly, however, they are aligned in our model (**Fig. S6F**), suggesting the possibility of a higher-order oligomeric bridge involving these cysteines and other copies of cofilin. We think it is a strong possibility that some other cross-linking molecule replaces fascin, and it would need to be able to interact directly with cofilin or with the new regions of actin exposed by cofilin binding.

If cofilactin bundles exclude fascin (**Fig. 1D & E**), what other actin binding proteins are regulated by this transition to cofilactin? It is known that cofilin inhibits binding of the Arp2/3 complex to actin (*58*), so one possibility is that the switch to cofilactin bundles prohibits the base of the filopodia from becoming integrated into the branched lamellipodial actin network, allowing it to move more freely. Indeed, our movies often show the base flexing back and forth quite freely within the growth cone body, or propagating waves generated by lateral movement of the tip (**Fig. 3E, Movie S5**). Myosin II function could also be regulated by cofilactin bundles. The pulling force of myosin II is known to account for about half of the retrograde flow in neuronal growth cones (*20*), and competitive inhibition between myosin II and cofilin has been demonstrated (*48*). This means that retrograde flow could be reduced in filopodia with a cofilactin bundle at their base, and this reduction in flow rate could account for the fact that filopodia with cofilin-rich regions were twice as long as those without one (**Fig. 1D**). Such an inverse correlation between retrograde flow and filopodial outgrowth (*19*) could, in part, be facilitated by the competing interactions of myosin II and cofilin with filopodial actin.

What does a transition region look like? Our models of fascin-linked actin bundles and cofilactin bundles provide a framework for exploring possibilities. Based on the models in Figure 4A, a completely abrupt transition between actin and cofilactin bundles would mean that 50% of the filaments within the bundle twist by 90° at the transition, perhaps severing individual filaments. If the transition is less abrupt, and spans a greater distance, it’s unlikely that all of the filaments would twist in a single cross-sectional plane, thus creating a broader inflection point (**Fig S4B)**. Based on our interfilament distances, the transition would also include a tighter packing of the filaments on the cofilactin side, but it’s not immediately clear how this would contribute to flexibility. Further tomography is needed to determine the structural mechanism of bending at these transition regions.

Do cofilactin bundles near the base represent a filopodial breakdown intermediate? They almost certainly do given cofilin’s documented severing function (*28*–*31*), and their prevalence near the base of filopodia, where turnover is known to occur (*20*). We do observe bundles collapsing and breaking within our movies, near the base of filopodia (**Fig. 3D, Movie S4)**, a process known to be regulated by non-muscle myosin II (*20*). Because of this and other studies showing synergy between cofilin and myosin (*47*), we believe cofilactin bundles function to regulate filopodial breakdown in conjunction with motor proteins. Current evidence suggests that cofilin severs actin filaments by generating intrafilament transitions between cofilactin and bare F-actin. The helical offset between these two portions of the filament is thought to disrupt intermolecular bonds within actin filaments to produce severing (*30*, *59*–*61*). Due to the competitive inhibition exhibited by myosin II and cofilin (*47*, *48*), myosin’s presence could result in actin filaments that are more sparsely decorated with cofilin, leading to more cofilactin/actin boundaries, and more filament severing.

Previous work in non-neuronal cells suggested that cofilin and fascin work synergistically to break down filopodial bundles (*43*). In their model, the twisting cofilactin filaments at the tip of filopodia compete with the fascin cross-linkers, leading to localized stress and increased severing events. In our model, fascin is ejected by the twisting and compressing of the filament bundle, leaving behind fully intact cofilactin bundles. Why cofilin and fascin would interact so differently across cell types is not immediately clear, but given the specialized nature of the growth cone, it could be that this mechanism for cofilactin bundles evolved under the specific pressures of neurite navigation.

Needless to say, there is still work to be done to fully understand the role of actin remodeling in filopodial structure and dynamics, as well as the interplay of cofilin and fascin with other actin binding proteins. These findings, however, point to a novel and complex role for cofilin in regulating filopodial structure and function that does not only include severing. Finally, given the antennae-like nature of filopodia in axon guidance, we hypothesize that the searching behavior afforded by cofilin-induced bending is essential for optimal space-searching, and, therefore, regulates the efficiency of targeted neurite outgrowth.

## Acknowledgments

We would like to thank Dr. Neal Waxham for insightful discussion and feedback throughout the writing of this manuscript, and Peter Swulius for help with custom Matlab scripts. We would also like to acknowledge support from the cryo-EM core facilities at the Penn State College of Medicine (Hershey, PA) and at the Penn State University Park (State College, PA) campuses.

## Materials and Methods

### Neuronal Cell Culture

Coverslips and EM grids were coated with 100 μg/mL poly-D-lysine (PDL) (Millipore) overnight. PDL was subsequently rinsed 3 times with Milli-Q H_2_O prior to cell plating. E18 Sprague Dawley rat hippocampi were acquired from BrainBits LLC and cultured according to their protocol. Cells were either cultured in NbActiv4 (BrainBits LLC) or Neurobasal (Gibco) plus 2% (v/v) B-27 Supplement (Gibco), both containing 1% Penicillin-Streptomycin (Gibco). For immunofluorescence experiments, cells were plated at 20,000 cells/cm^2^ on 12 mm German glass coverslips (Electron Microscopy Sciences) (either #1 or #1.5 coverslips were used, depending on the microscope). For cryo-ET, cells were plated at 30,000 cells/cm^2^ on 200 mesh gold R 2/2 carbon Quantifoil EM grids (Electron Microscopy Sciences).

### Cell Vitrification

On DIV 1, 2, or 3 (∼24-72 hours after cell plating), cells were vitrified in liquid ethane using a Vitrobot Mark IV (ThermoFisher Scientific). Prior to vitrification, 10 nm gold fiducial markers (Ted Pella) coated in 1% BSA were diluted 1:4 or 5 in conditioned culture media and 3μl of this mixture was added on top of each grid. Grids were then blotted by hand from behind for ∼2 seconds with Whatman filter paper and immediately plunge-frozen in liquid ethane.

### Cryo-ET

EM was performed on a ThermoFisher Scientific FEI Titan Krios G3i 300kV FEG TEM equipped with either a 4k × 4k pixel K2 or 6k × 4k pixel K3 Summit direct electron detector (Gatan). A GIF energy filter (Gatan) with a slit width of 20kV was used during operation and images were collected in electron counting mode. Magnification was typically 26,000×, corresponding to a pixel size of 4.306 Å on the K2 detector and 3.326 Å on the K3. Defocus was −6 to −8 μm. Each tilt series was collected from −60° to +60° with tilt increments of 1° or 2° between images, generally in a bidirectional tilt scheme. A total electron dose of ∼150 electrons/Å^2^ was used for each tilt series.

### 3D Tomographic Reconstruction, Tomographic Analysis, and Subtomogram Averaging

Tomogram alignment and reconstruction were performed in the IMOD software package (*62*). Alignment was executed using a fiducial model if possible; otherwise 2 iterations of patch tracking were used. Reconstruction was done by weighted back-projection. All tomograms shown in the figures of this manuscript were filtered for higher contrast either with a median filter (kernel size of 3 pixels) using the clip function in IMOD or a SIRT-like filter, also in IMOD.

Cofilactin twist-lengths were measured in IMOD and calculated using multiple twists (usually 5-10 per filament; 56 measurements total) on a single filament simultaneously. In total, 9 filaments from 3 bundles were measured.

Dynamo was used for subtomogram averaging of individual actin and cofilactin filaments (as in **Fig. 2D-E**). For cofilactin averages, particles were placed every 27.6 Å and were rotated −162.1° (the approximate axial rise and twist of each actin monomer in cofilactin (*37*–*39*). For actin averages, particles were placed every 27.6 Å and were rotated −166.6° (*63*). 3,963 particles were used for cofilactin averages and 1,603 particles were used for actin averages. For subtomogram averaging of filament pairs (as in **Fig. 2B-C**), the software package PEET was used. Rigid body fitting of atomic structures to our EM maps was done in Chimera using the fitmap command.

### Neural Network Segmentation, Filament Centerline Extraction, and Nearest Neighbor Analysis of Tomograms

Segmentations were generated using Dragonfly software, Version 2021.1 for Windows (Object Research Systems (ORS) Inc, Montreal, Canada, 2020; software available at www.theobjects.com/dragonfly). Full resolution (4k × 4k or 6k × 4k) tomograms were loaded into Dragonfly and filtered with a histogram equalization filter followed by a 3D Gaussian smoothing filter to boost signal. A small rectangular box containing the feature of interest was selected out of the full tomogram. All voxels within this box were hand segmented as either the feature of interest or negative data. The box was used as a training mask for training a neural network. Using the Deep Learning Tool in Dragonfly, a multi-slice (5 slices) U-Net was generated for a two-class semantic segmentation. The multiROI generated from hand segmenting the voxels was used as the training output, the filtered tomogram was assigned as the training input and the mask ROI was used to limit the training to the segmented region. All segmentations required some manual clean-up after AI segmentation. The segmented feature of interest was then exported as a binary TIFF.

Binary images were converted from TIFFs to MRC image stacks using the tif2mrc function in IMOD and were imported into Amira where filament centerlines were extracted using its filament tracing tool (*64*). A minimum interfilament distance of 8 nm was assigned, as this is the approximate width of an actin filament, therefore, two filaments centerlines could not be any closer than this. The outputted data possessed coordinates of points along filaments from which nearest neighbor distances were calculated in MATLAB (MathWorks) using a custom script. Nearest neighbor distances greater than 3 standard deviations above and below the mean were eliminated as outliers.

The transition region tomogram in Figure 3G was produced using a 4×-binned (1k × 1k pixel) tomogram. As above, contrast was enhanced with a histogram equalization filter followed by a 3D Gaussian smoothing filter. Filaments were then hand segmented in Dragonfly.

### Modeling

For figure S6, UCSF Chimera (www.cgl.ucsf.edu/chimera/) was used to build a seven-filament unit of hexagonally packed filaments, using both F-actin (PDB ID: 3G37) and cofilactin (PDB ID: 3J0S) atomic models. Filaments were placed at distances from one another based on our interfilament distance measurements (12.3 for fascin-linked bundles and 11.5 for cofilactin bundles). The atomic model for fascin (PDB ID: 3P53) was fit in by hand, based on modeling done by Aramaki et al (*22*). For cofilactin bundles, filaments were rotated with respect to each other, based on our observations within raw tomograms (**Fig. 4A & B**).

### Fixation and Immunofluorescence

If they were to be stained with phalloidin for actin staining, neurons were fixed on DIV 1 with a 4% PFA solution (1-part 16% PFA-Electron Microscopy Sciences, 1-part Milli-Q H_2_O, 2-parts 2× PBS with 8% sucrose) at room temperature for 10 minutes. This was followed by a 5-minute rinse with 50mM glycine in 1× PBS and 3 brief rinses in 1× PBS. Cells were then permeabilized by acetone (pre-chilled to −20°C) for 1 minute, followed by a final 1-minute rinse in 1× PBS prior to addition of blocking buffer. The cell in Figure S1A was permeabilized with 0.5% Triton-X 100 for 10 minutes prior to the addition of blocking buffer. If fascin was being labeled instead of actin, cells were simultaneously fixed and permeabilized with methanol (pre-chilled to −20°C) in the −20°C freezer for 20 minutes. This was followed by 3 brief 1× PBS rinses as above. See the Supplementary Text and Figure S1 for more details.

Blocking buffer (2% normal goat serum and 1% w/v BSA in 1× PBS) was added for 15 minutes prior to antibody labeling. Primary antibodies (diluted in blocking buffer from above, 30-minute incubation at room temperature): rabbit anti-cofilin at a 1:1000 dilution (Sigma-Aldrich), mouse anti-fascin at a 1:500 dilution (Santa Cruz Biotechnology). Secondary antibodies (also diluted in blocking buffer from above, 30-minute incubation at room temperature): goat anti-rabbit conjugated to Alexa Fluor 594 at a 1:500 dilution (Abcam), goat anti-mouse conjugated to Alexa Fluor 488 at a 1:500 dilution (Abcam). Primary and secondary antibodies were rinsed 3 times for 5 minutes per rinse with a rinsing buffer (blocking buffer from above diluted 1:10 in 1× PBS). If phalloidin (conjugated to Alexa Fluor 488, Invitrogen) was used, it was diluted 1:66 in the blocking buffer and added after all other antibody labeling and rinsing steps. Afterwards, phalloidin was rinsed 3 times in the rinsing buffer for only ∼30 seconds per rinse. All coverslips were rinsed once with 1× PBS and once with Milli-Q H_2_O prior to being mounted on a slide with a hardset mounting medium containing DAPI (Biotium) and allowed to cure overnight in the dark at room temperature.

### Fluorescent Imaging

Images of growth cones where actin and cofilin were labeled were taken on a DeltaVision Elite Deconvolution widefield microscope equipped with a cooled EM-CCD. Images were taken on either a 100× or 60× oil objective with DAPI, FITC (for Alexa Fluor 488 visualization), and Texas Red (for Alexa Fluor 594 visualization) filters. Growth cones where fascin and cofilin were labeled were imaged on a Zeiss Axio Observer equipped with a Colibri 7 light source. Here a 100× oil objective was used with a 110 HE LED filter set and EGFP, mRF12, and DAPI filter settings active. Z-stack images were taken with the automated “optimal” Z-section distance used. Additionally, the Apotome 2 was used for optical sectioning of the images. Raw Apotome images were deconvolved using Zeiss’ Zen Blue software.

### Analysis of Fluorescent Images

Brightness and contrast of images were adjusted using FIJI (https://imagej.net/software/fiji/). Additionally, FIJI was used for measurements of bundle and filopodial lengths (**Fig. 1D**) as well as the quantification of the frequency of cofilin-rich bundles in growth cones. All line scan intensity profile measurements were made in Zeiss’ Zen Blue software on a central slice from each Z-stack, where the fluorescence intensity was brightest. For the measurements made along the long axis of filopodia (**Fig. 1E**), one end of the line was placed at the bottom of the cofilactin portion of filopodia and the other end was placed at their most distal tip, as visualized by fascin staining. A correlation coefficient was then calculated from the resulting intensity line graphs of each filopodium measured.

Colocalization analysis was also performed in the Zen Blue software. To determine an optimum threshold for colocalization analysis, three random growth cones were outlined as regions of interest and the “Costes” function within Zen Blue was used before an average threshold pixel value for each channel was calculated. Then, whole filopodia, or just the “transition region” (defined in the Supplementary Text), were boxed out. The Pearson’s Correlation Coefficient was calculated based on the above threshold values.

### Nucleofection

Primary E18 rat hippocampal neurons (BrainBits LLC) were dissociated from hippocampal tissue slices according to the company’s protocol. After dissociation, cells were nucleofected with a Nucleofector 2b (Lonza) using the company’s protocol for rat neuronal cells. Briefly, 1 to 2 million cells were placed in 100 μL of nucleofection solution with a total of 3 μg of plasmid DNA (when two plasmids were used, 1.5 μg of each plasmid was added). Cells were placed in the nucleofector unit and nucleofected using program G-013. After transfection, cells were plated on PDL-coated, 35 mm, glass-bottom dishes (MatTek). Here PDL was the same concentration as above, but plates were rinsed just once with Milli-Q H_2_O and dried in a cell culture hood for ∼1 hour prior to use. Cells were incubated overnight in Neurobasal (Gibco) with 10% FBS (R&D Systems) and 1% Penicillin-Streptomycin (Gibco). After overnight incubation, the media was changed to Neurobasal media with 2% (v/v) B-27 Supplement (Gibco) instead of FBS.

### Live Cell Imaging

All movies were made on a Zeiss LSM980 using the Airyscan 2 detector and a 40× dipping objective at room temperature. Excitation laser settings were as follows: tdTomato: 561 nm and EGFP: 488 nm. All movies collected were over a 10-minute timescale where images were taken every 3 seconds. Movies were processed using the Airyscan image processing tool in Zen Blue with default settings.

### Quantification of Live Cell Imaging

Maximum intensity projections (MIPs) were made from images across 40 time points (2 minutes-worth) using FIJI. The width of filopodia was then measured in three locations: the widest point of the cofilin-labeled portion, the widest point of the actin-labeled portion (more distal than cofilin), and the point at which the two signals met (the inflection point/transition region as described above, in the Main Text, and in the Supplementary Text). The width in this case represented the amount that each filopodial region moved from side-to-side during the 2-minute time block. “Searching” filopodia were defined as those whose actin width was at least 2-fold greater than their cofilin width in the MIPs.

### Plasmids

All plasmids were acquired from Addgene and verified by sequencing using the associated primers on the Addgene website. The following plasmids were used: pEGFP-N1 human cofilin (Addgene 50859) and td-Tomato-LifeAct 7 (Addgene 54528).

## Supplementary Text

### Permeabilization Methods for Immunofluorescence of Growth Cone Cofilactin Bundles (Figs. 1 and S1)

Figure 1 and S1 contain immunofluorescence microscopy images that show cofilactin bundles in growth cones. Depending on the additional staining performed (actin or fascin), different fixation and permeabilization protocols were used.

We have observed that the use of Triton-X 100 as a permeabilizing agent before immunolabeling disrupts cofilactin bundle staining (**Fig. S1A**). Cofilin is still labeled throughout the cell, but there are no discernable aggregates of cofilin signal beneath filopodia. In contrast, the use of organic solvents (ice-cold acetone or methanol) for permeabilization preserves cofilactin bundle staining. Interestingly, this phenomenon has also been shown for cofilin-actin rods.

When staining cofilin and actin, acetone was used as a permeabilizing agent as methanol destroys the phalloidin binding site on actin (**Fig. S1B**). When staining cofilin and fascin, methanol was used as a fixative instead of PFA (**Fig. S1C**). Methanol is used since PFA disrupts the binding of our fascin antibody (data not shown). Methanol acts as a fixative and permeabilizer in this case. Cofilactin bundles also stain well with PFA fixation and methanol permeabilization (data not shown). We have noticed that acetone slightly diminishes phalloidin and cofilin signal and that cofilin bundle morphology is best preserved with methanol fixation and permeabilization. This is why filopodial bundle length and cofilactin bundle frequency measurements were made on growth cones treated in this way.

### Definition of the Filopodial “Transition Region” (**Fig. S2**)

For colocalization analysis (**Fig. S2**), either an entire filopodium was measured, or, if measuring the transition region, a rectangular box that ran 400-nm along the length of the filopodia and was equal to their width was placed in the middle of the filopodia where, on the merged image with both channels activated, the fascin and cofilin signals met. The box was placed immediately distal (towards the fascin-labelled portion) to this spot. This location is diagrammed in Figure 1E. The transition region is also referred to as an inflection point throughout this manuscript.

---

**Fig. S1.**
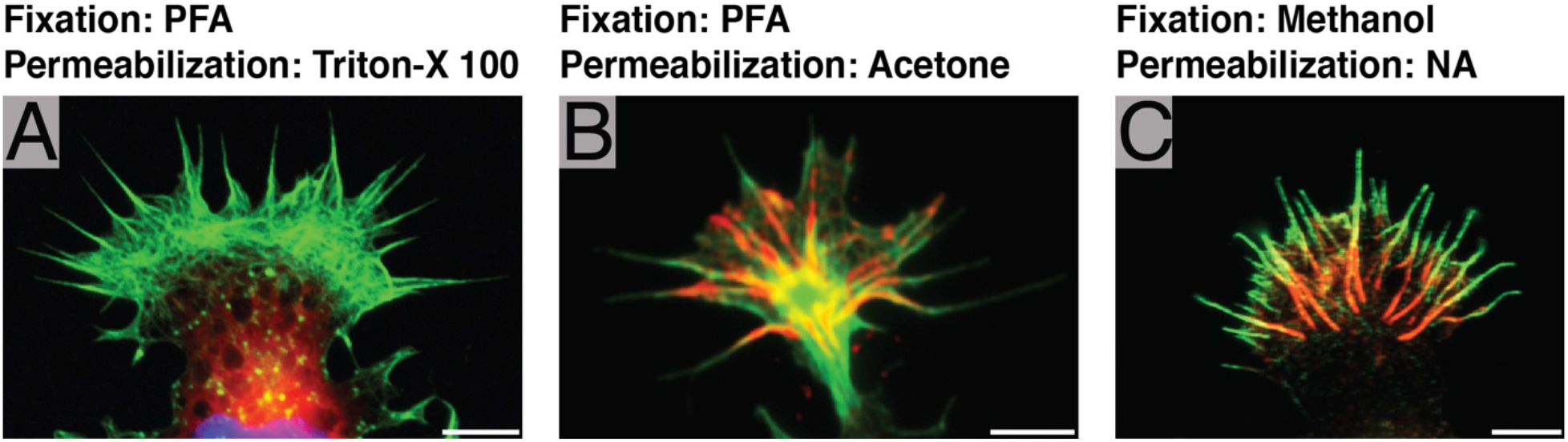
Effect of permeabilization on cofilactin bundle visualization. As discussed in the Supplementary Text, multiple fixation and permeabilization protocols were used prior to immunolabeling growth cones. In general, permeabilization with Triton-X 100 seems to prohibit cofilactin bundle labeling, while permeabilization with organic solvents does not. In (A) and (B), green is actin (phalloidin) and red is cofilin. In (C) green is fascin and red is cofilin. **(A)** Fixation with 4% PFA followed by permeabilization with 0.5% Triton-X 100 enables cofilin labeling, but no cofilactin bundles are visible. **(B)** Fixation with 4% PFA and permeabilization with ice-cold acetone preserves cofilactin bundle morphology, although slightly reduces phalloidin and cofilin signal intensity. **(C)** Fixation with methanol preserves cofilactin bundle morphology but destroys phalloidin’s binding epitope, so fascin is used to label the more distal filopodial regions. Methanol also acts as a permeabilizing agent. Scale bars are all 5 μm.

**Fig. S2.**
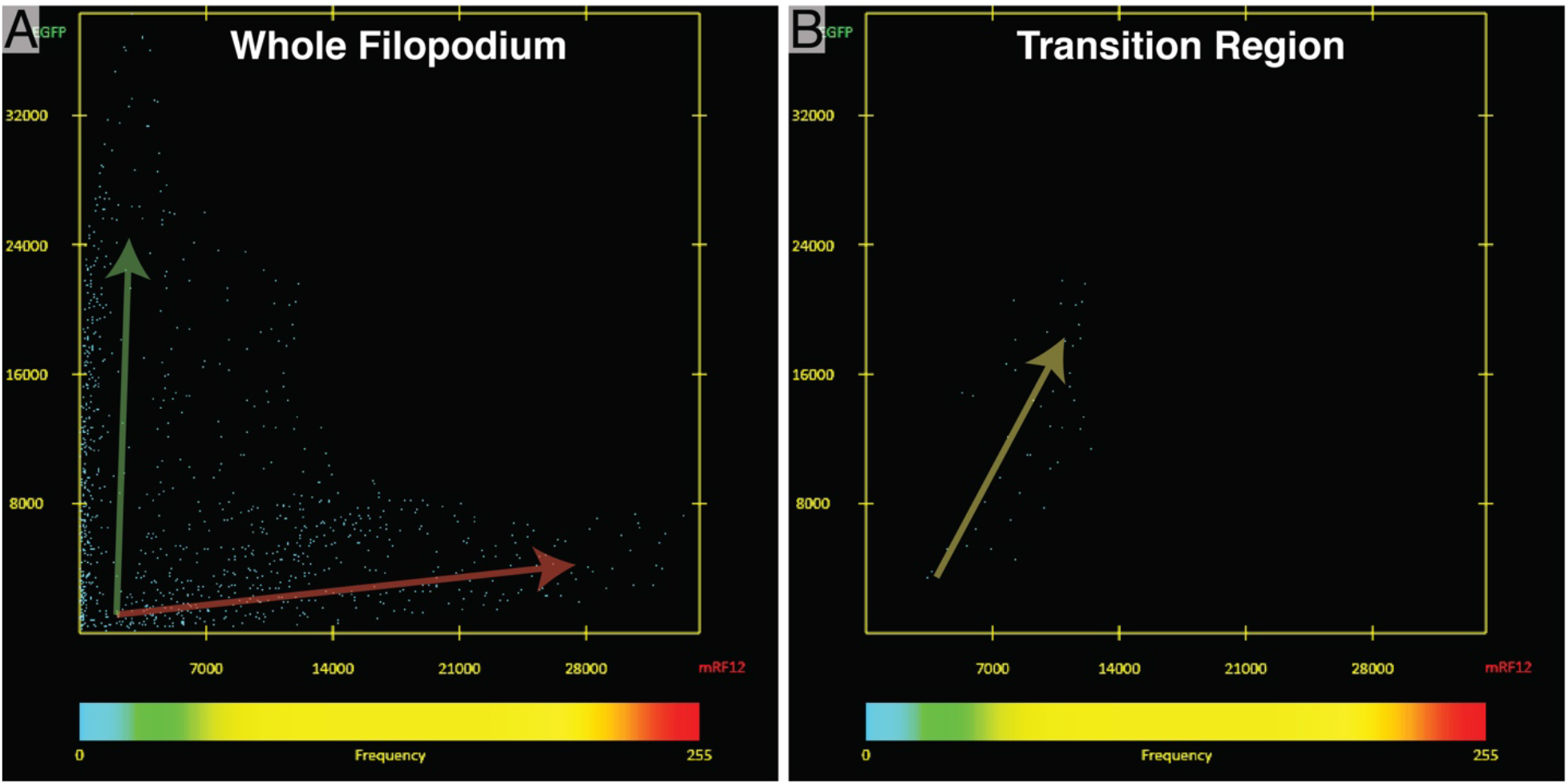
Colocalization of fascin and cofilin in neuronal growth cone filopodia. **(A) & (B)** Representative scattergrams resulting from colocalization analysis of fascin and cofilin in a whole filopodium (A) and the transition region (B) (as defined in the Supplementary Text and shown in Figure 1E). The average Pearson’s Correlation Coefficient for whole filopodia was −0.28 +/− 0.19 (+/− S.D.) and for the transition region was 0.39 +/− 0.29. Green and red arrows in (A) show pixels that predominately had fascin or cofilin signal, respectively. The yellow arrow in (B) brings attention to the linear relationship between pixel intensities in the green and red channels in the transition region.

**Fig. S3.**
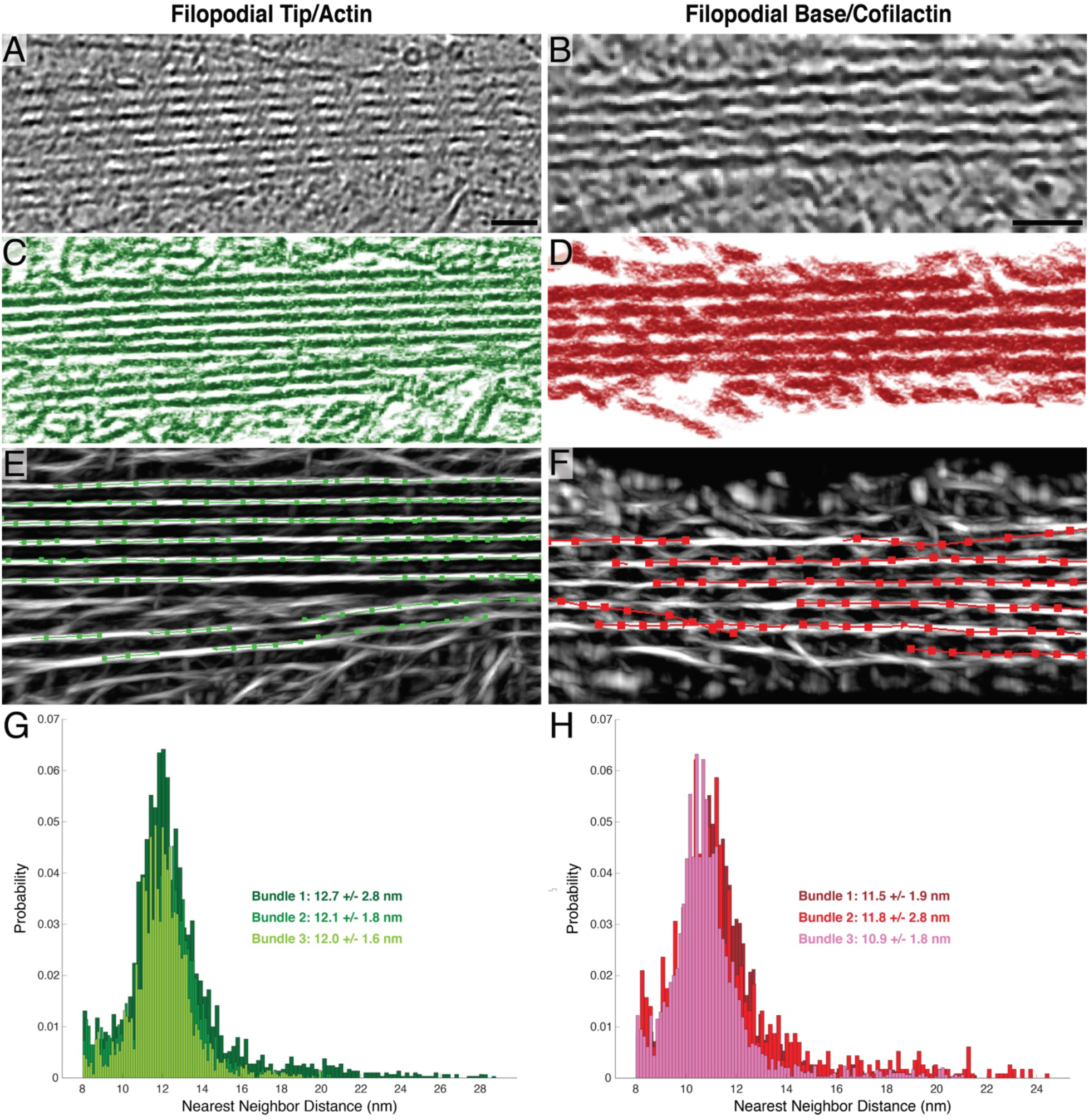
Neural network segmentation and interfilament distance analysis workflow. **(A-F)** The workflow for the segmentation and filament centerline extraction is displayed in the following way (filopodial tips/actin bundles and filopodial bases/cofilactin bundles were analyzed the same as one another): (A) and (B) show example raw tomograms that were segmented. (C) and (D) show annotated actin and cofilactin filaments, respectively, after segmentation in Dragonfly. (E) and (F) show centerlines that were extracted from filaments using Amira. The points along each line were the points from which nearest neighbor calculations were made in MATLAB using a custom script. **(G & H)** Three bundles of each type were segmented and analyzed. Nearest neighbor measurements are plotted as histograms and all bundles from each type are overlaid together. Bundles from each type overlap heavily, indicating a consistent structure across multiple filopodia. Scale bars in (A) and (B) are 40 nm.

**Fig. S4.**
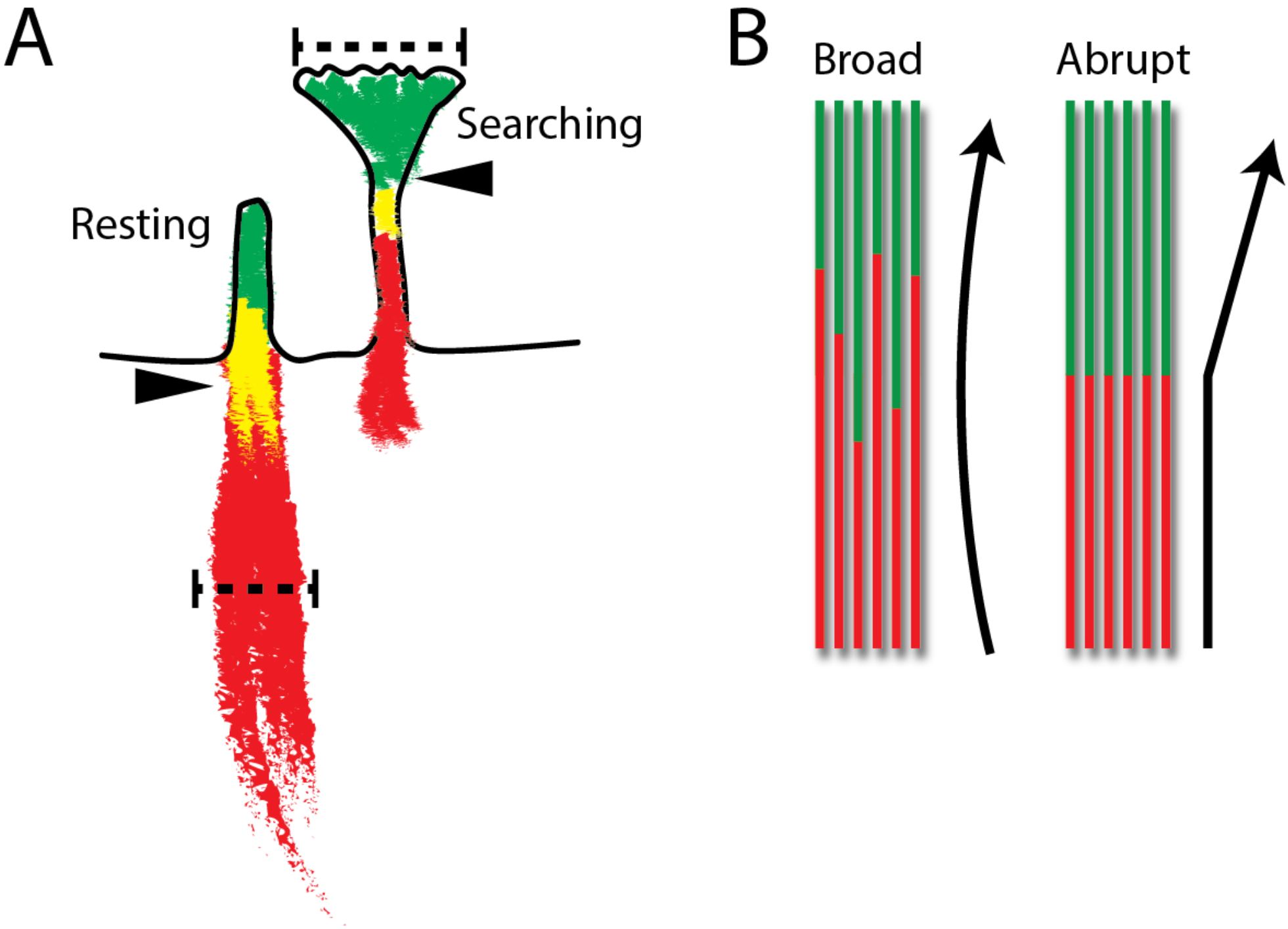
Proposed impact of the transition region on growth cone filopodial behavior. **(A)** Schematic representing the signal distribution in maximum intensity projections (MIPs) of neuronal growth cone filopodia, as shown in Figure 3B and C. The width of each region was measured as shown here. The cofilin (red), inflection point/transition region (yellow), and actin (green) signal widths were measured at their widest point in MIPs made from 2 minutes-worth of images. Arrowheads point to the inflection point at the regions of colocalization between actin and cofilin **(B)** Schematic showing how the distribution of actin-to-cofilactin transitions could impact the bending angle of filopodia such as those in A. In one, the location of the transition is more broadly defined, and any cross-section through it would show a mixture of cofilactin and fascin-linked normal actin. In the other, the transition is more abrupt, and the shift from normal actin to cofilactin occurs all at once. We hypothesize that the transition region of resting filopodia is more broad and closer to the lamellipodial veil. This results in subtle translations of the filopodia, as shown by the left arrow in (B), and most movement occurs in the cofilactin base. Conversely, we hypothesize that searching filopodia have more abrupt transition regions that are more distal than the lamellipodial veil. This results in more sharp angular translations of the filopodial tip, as shown by the right arrow in (B).

**Fig. S5.**
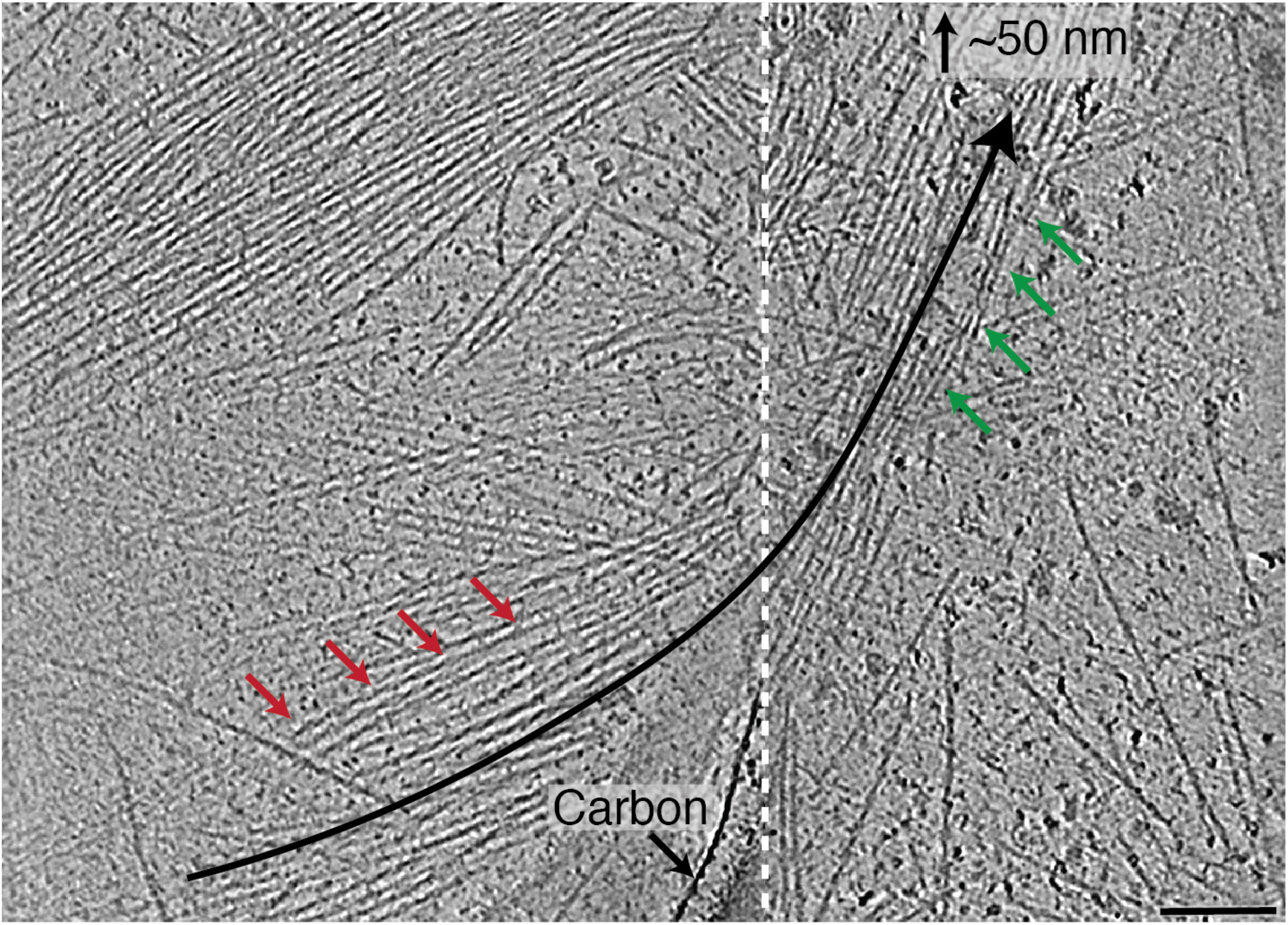
Example of a prospective transition region in a growth cone filopodium. This filopodium possesses a fascin-linked filopodial bundle (right, green arrows) that transitions into a cofilactin bundle (left, red arrows). The left and right sides of the tomogram are separated with a white dashed line because the normal actin bundle is on the carbon surface of the EM grid, while the cofilactin bundle is in a hole, and therefore they are at very different Z-heights. The black arrow follows the bundle and illustrates how the two regions are oriented at substantially different angles. Scale bar is 100 nm.

**Fig. S6.**
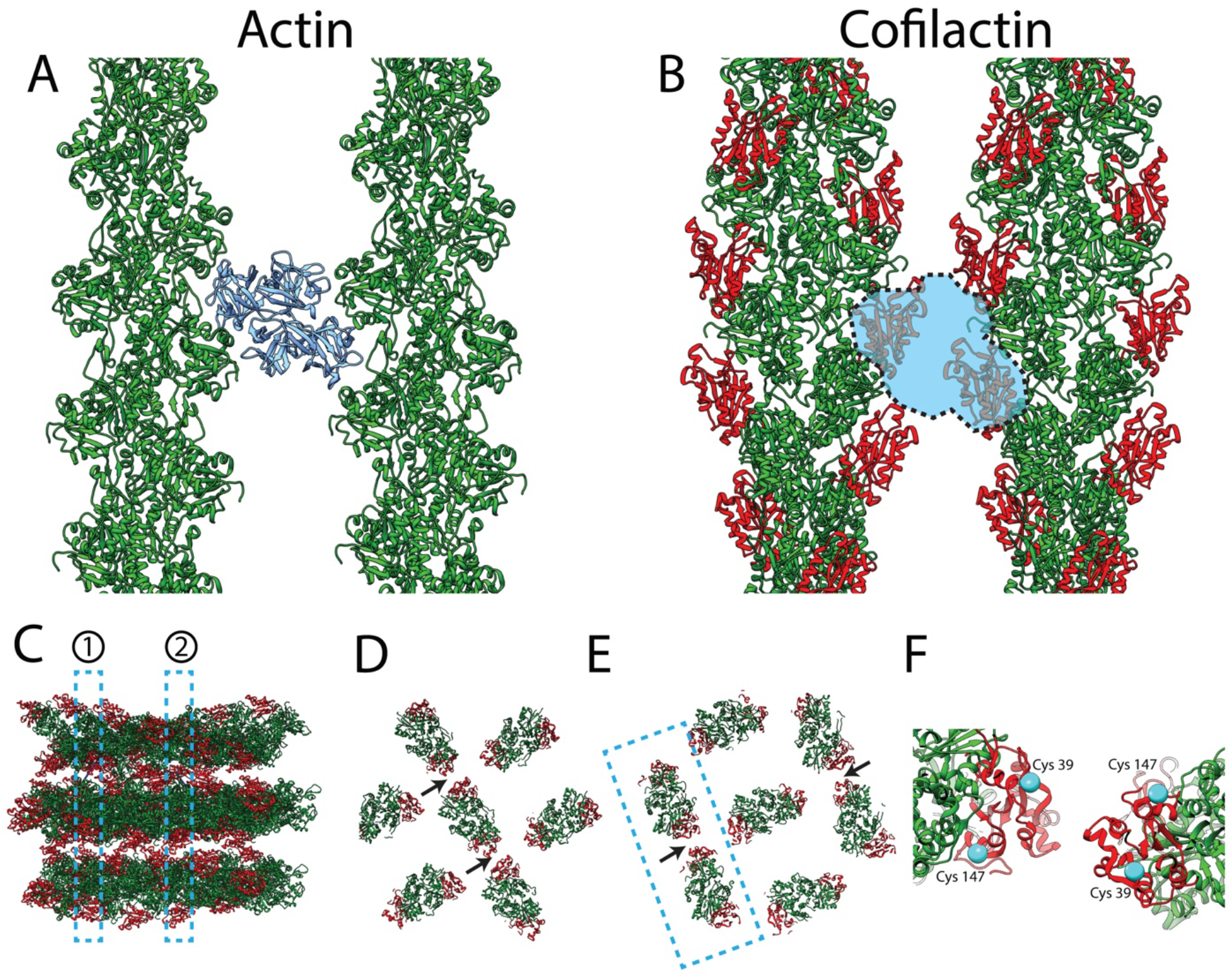
Potential cofilactin interactions. **(A & B)** Top-down views of fascin-linked actin (A) and cofilactin (B) filaments. Both pairs of filaments are shown at interfaces where they are in phase with one another. In (A), fascin (blue) is shown in its actin-binding pockets (*22*), two sites that are sterically blocked by the presence of cofilin in our model of cofilactin bundles (B). F-actin is green and cofilin monomers are red. **(C)** Sideview of model cofilactin hexagonal unit. Dashed regions labeled 1 and 2 illustrate cross-sections shown in (D) and (E), respectively. F-actin is green and cofilin is red. **(D & E)** Cross-sections through different regions of the hexagonal unit shown in (C). Arrows point to the place where cofilin monomers on neighboring filaments are closest to each other (∼2 nm apart at their closest residues). **(F)** Zoomed-in view of neighboring cofilins like those in the boxed-out region in (E). Cys39 and Cys147 are displayed on each (cyan).

Movie S1. **Cryo-tomogram of a growth cone filopodium.** Tomographic reconstruction of a growth cone filopodial protrusion. Tightly bundled actin filaments nearly fill the entire cytoplasmic space, and fascin cross-linking densities can be seen at regular intervals along the bundle.
Movie S2. **Cryo-tomogram near a growth cone filopodium’s base.** Tomographic reconstruction of a growth cone filopodial base. Individual actin filaments surround a bundle of hyper-twisted cofilactin filaments.
Movie S3. **Movie of whole cell co-expressing fluorescent cofilin and LifeAct.** Neurons were transfected with pEGFP cofilin (red) and tdTomato Lifeact (green). The movie is pseudo-colored such that Lifeact is green and cofilin is red to match the color scheme used throughout this study. Images were taken every 3 seconds for 10 minutes. This is the same cell as in Figure 3A.
Movie S4. **Breakage of filopodium in the cofilactin bundle region.** Zoomed in movie of filopodia depicted in figure 3D.
Movie S5. **Cofilactin bundle “wave” during filopodial motion.** Zoomed in movie of filopodia depicted in figure 3E.
Movie S6. **Movie of filopodial bend.** Zoomed in movie of filopodia depicted in figure 3F.
Movie S7. **Cryo-tomogram at a bend in a filopodium.** Tomographic reconstruction of the region depicted in figure 3G.

## Notes

### Competing Interest Statement

The authors have declared no competing interest.

### Summary of Updates

Added new supplemental figures and a revised abstract. We also updated a citation that was in the incorrect format.

